# Identification and structural characterization of pseudogenes in *Fusarium graminearum*

**DOI:** 10.1101/2024.09.17.613474

**Authors:** Domenico Rau, Chiara Maria Posadinu, Maria Leonarda Murgia, Davide Fois, Andrea Porceddu

**Affiliations:** Dipartimento di Agraria, Università degli Studi di Sassari, Viale Italia 39/A 07100 Sassari

**Keywords:** Pseudogene, Pseudogene classification, Intron-exon structure, Pseudogene Identification, Duplication mode, Gene ontology

## Abstract

Pseudogenes provide valuable insight into the evolutionary history of genomes which can be challenging to ascertain through the examination of functional loci alone. This study presents the findings of a comprehensive, two-step genome-wide survey for the identification and characterization of pseudogenes in *Fusarium graminearum,* the primary causal agent of wheat head blight.

By analysing the sequence homology between non-coding regions of the genome and predicted protein sequences, we identified regions with homology to putative paralogous functional sequences. These regions were characterised in terms of their matching sequence structure and position. Most identified pseudogenes were mapped within the fast-evolving genomic compartment and were derived from transposition events. The number of processed and putatively retroposed pseudogenes was found to be comparable. The number of identified pseudogenes was low, which is consistent with the low number of gene duplicates in *F. graminearum*. No compelling evidence was found to suggest that pseudogene formation can be explained by evolutionary accidents during gene family expansion or as caused by RIP-associated mutagenic events. Notably, about one-third (144/436) of the pseudogenes were found to overlap with untranslated or intron sequences of functional loci, indicating the potential to be transcribed. Using *Fusarium* comparative genomics, we identified genomic regions with homology to genes lacking functional orthologs in *F. graminearum*. These were investigated as putative unitary pseudogenes and, in some cases, their original functions were completely lost after the radiation of *F. graminearum*.

Interestingly, the paters of 18 loss-of-function pseudogenes showed homology to domains previously identified in proteins involved in pathogenesis.

## Introduction

Pseudogenes are copies of genes that have lost the ability to encode functional proteins as a consequence of disabling mutations. They are also referred to as genomic fossils and have been considered “junk DNA” (Balakirev and Ayala, 2003; Tutar, 2012; Vanin, 1985). Despite their widespread acceptance and utilisation, these definitions are increasingly challenged by recent research (Guo et al., 2009; Pink et al., 2011). These have demonstrated that pseudogenes can be expressed and interfere with the function of cognate paralogous genes via a variety of mechanisms (Cheetham et al., 2019). The regulation of gene function may entail post-transcriptional gene silencing mechanisms or may act through the residual activity of regulatory sequences that have been retained in the pseudogene sequence (Cheetham et al., 2019; Poliseno et al., 2015; Salmena et al., 2011).

Even pseudogenes with an as-yet undefined biological role may prove highly informative, as they often reveal aspects of genome evolution more effectively than functional loci (Cheetham et al., 2019; Sisu et al., 2014). For example, the study of pseudogenes can assist in identifying genes that have been inactivated or lost during pathogen evolution, thereby shedding light on the genes and pathways that are critical for pathogenicity and/or virulence (Feng et al., 2022; van der Burgt et al., 2014; Yang et al., 2024). In a plant patho-system, pseudogenes may play a role in the evolutionary arms race between pathogens and their host plants. The selective pressure exerted by host plants may lead to the inactivation of effectors that correspond to the receptor-like protein encoded by the resistance gene (Feng et al., 2022; Yang et al., 2024). Conversely, pseudogenes may serve as a reservoir of genetic material that can be reactivated or repurposed to contribute to pathogenicity. Comparative studies have shown that the formation of pseudogenes in pathogenic fungi with different host plants or lifestyles can reflect the evolutionary history of the pathogen past (De Wit et al., 2009; Goodwin et al., 2011). The genes encoding secreted proteins, including proteases, were observed to undergo pseudogenisation more frequently in *C. fulvium* than in *D. septosporum* (van der Burgt et al., 2014). Most of the pseudogenes identified in these two closely related fungi are not shared, suggesting that they are associated with adaptation to a different host (tomato *versus* pine) and lifestyle (biotrophic *versus* hemibiotrophic) (De Wit et al. 2009).

Based on the intron-exon structure, pseudogenes are typically classified into two main types: non-processed (or duplicated) and processed (or retroposed) (Mascagni et al., 2021; Zhang et al., 2006). Non-processed pseudogenes originate from genome or chromosome duplication and typically retain the intron-exon structure of their pater loci. Processed pseudogenes are genomic copies of cDNA reverse transcribed from messenger RNA (Baertsch et al., 2008; Tan et al., 2016). These pseudogenes are usually intron-less and exhibit a stretch of adenines at their 3’ end (Baertsch et al., 2008). An alternative mechanism for the generation of pseudogenes is through ectopic duplications, whereby genomic sequences are copied to ectopic (‘out of place’) sites during a process of single-strand break repair involving Synthesis-Dependent Strand Annealing (SDSA) (Buchmann et al., 2012; Mascagni et al., 2021; Wicker et al., 2010). Pseudogenes generated by this latter mechanism rarely cover the full length of the pater locus (Mascagni et al., 2021; Prade et al., 2018). Different taxa showed different frequencies of pseudogene types. For instance, processed pseudogenes are prevalent in mammals, whereas plant genomes exhibit a low ratio of processed *to* non-processed pseudogenes (Takuno et al., 2008; Xu et al., 2008). The location and structure of pseudogenes in relation to the functional paralogous sequence may also have relevant implications for gene evolution. Pseudogenes with conserved gene structure and mapping near their pater loci have been identified as implicated in gene conversion events (Takuno et al., 2008; Xu et al., 2008). Furthermore, an examination of pseudogenes in both wild and domesticated barley accessions has demonstrated their capacity to contribute to sequence diversity and evolution. In some cases, they can resurrect or acquire new functions, thus representing a reservoir of genic-like sequences (Prade et al., 2018).

In species that have undergone few genomic or segmental duplications, most genes are present as single copies (singletons). Consequently, their eventual inactivation by a disabling mutation result in the loss of gene function (Mitchell and Graur, 2005; Zhang et al., 2010). These pseudogenes are defined as ‘unitary’ to indicate that they devoid of any functional paralogous sequences (Harrison et al., 2003; Zhang and Gerstein, 2004; Zhang et al., 2010). The analysis of unitary elucidated the evolutionary trajectories of closely related species and in some cases have been instrumental in explaining adaptive processes more than extant biological function (Sisu et al., 2014; Zhang et al., 2010).

The genus *Fusarium* is of primary relevance within the Ascomycetes, comprising many species that are pathogenic to crops. Comparative genomics has shown that *Fusarium* species exhibit a genome architecture comprising two distinct compartments. The first compartment is highly conserved, displaying a high degree of gene collinearity between Fusarium species, while the second compartment shows a lower degree of gene collinearity and is enriched for species-specific genes and genes involved in the synthesis of secondary metabolites important for virulence (Cuomo et al., 2007; Dong et al., 2015; Torres et al., 2020).

Some fungi have evolved a defense mechanism against repeat elements, designated RIP (Repeat Induced Point Mutations), which introduces C to T transitions in duplicated sequences (Galagan and Selker, 2004). Theoretical calculations have demonstrated that RIP would result in the inactivation of a gene through the introduction of a stop codon within a few generations (Hane and Oliver, 2008; van Wyk et al., 2021). Although RIP is induced by duplicated sequences exceeding 400 bp, once initiated, it has the potential to induce point mutations beyond the duplicated sites, thus involving neighboring sequences. It is therefore expected that signatures of RIP activity and pseudogene abundance will be associated, which would have implications for genome structure and function. Most of the available information on the mechanisms leading to collinearity loss has been obtained from the analysis of functional genes, with relatively little knowledge derived from genome-wide analysis of pseudogenes. While the occurrence of RIP has been experimentally demonstrated in multiple Fusarium spp (Cuomo et al., 2007; Galagan and Selker, 2004), a comprehensive analysis of the relationship between RIP occurrence, gene decay and loss of collinearity remain lacking.

*F. graminearum*, the major cause of head blight, is regarded as the Ascomycetes with the most accurate genome annotation to date (Cuomo et al., 2007; King et al., 2017; Lu et al., 2022). Long-read sequencing of the transcriptome has revealed that approximately 60% of the 17,000 loci can produce alternative isoforms (Lu et al., 2022). Detailed transcriptome analyses have revealed that most of the alternative splicing (AS) events are due to intron retention which resulted in altered open reading frame (ORF) length and, consequently, in an increase in the complexity of the *F. graminearum* proteome (Lu et al., 2022). Moreover, Qi et al. (2024) have recently reported a detailed analysis of loci with premature stop codons which can be subjected to restorative RNA editing in *F.graminearum*. Several of these loci played crucial roles in germ tissue development and RNA editing was found crucial for their functioning. However, knowledge of pseudogenes in intergenes and non coding YL regions is still missing. In the present study we refine the annotation of the *F. graminearum* genome by applying a pipeline for pseudogene identification in non-coding genomic regions. We integrate the annotation of F. *graminearum* pseudogenes with (Qu) a curated annotation of pseudogenes in non-coding regions of YL genome with homology to paralogous loci and a comprehensive list of unitary pseudogenes. All these data are integrated with the YL-1 annotation and will be useful to analyse the possible involvement of pseudogenes in transcriptome evolution and/or regulation.

## Results

A total of 1862 pseudogenes matching 248 functional loci (hereafter defined as ‘paters’) were identified in non-coding regions of the *F. graminearum* YL genome (Supplementary Table S1). Two-hundred and seven (83.46 %) pater loci matched a single pseudogene. Of the remaining 41, only 25 matched more than two pseudogenes (Supplementary Table S1 for details).

A total of 1426 pseudogenes were identified in the sub-telomeric region of chromosome 4, forming a repeated array of seven sequences that matched a cluster of seven paters (FG4G31040, FG4G37170, FG4G37180, FG4G37190, FG4G37230, FG4G37240, and FG4G37250). The pseudogenes in the arrays showed similar features suggesting a single pseudogenisation event that preceded the duplications. Consequently, these pseudogenes were excluded from further analysis.

### The copy number of pseudogenes in *F. graminearum* is not indicative of past gene family expansion

Most of the pseudogenes were unambiguously assigned to a pater locus. Of the 436 pseudogenes identified, 352 (80.73%) matched to a single functional locus, while 58 (13.30 %) and 26 (5.96%) matched to two or more functional homologous loci, respectively (Tab. S1). In these two last cases, the pater assignment was resolved based on the highest Exonerate alignment score (see Supplementary Table S2).

We investigated the relationship between the number of duplications of the paters (hereafter defoned as Duplication Depth, DD) and the number of pseudogenes they aligned with. Most of (182 of 248) *F. graminearum* paters lacked any duplicated copies and were therefore classified as singletons (Tab. S1). No significant association was observed between the DD of the paters and the number of pseudogenes identified (χ^2^=2,501; P = 0.138; Supplementary Table S3). Of the 181 singleton paters (DD = 1), 156 matched a single pseudogene and 25 matched more than one pseudogene. Of the remaining 66 non-singleton paters (DD > 1), 51 matched a single pseudogene and 12 matched more than one pseudogene (Tab. S1).

### Most of *F.graminearum* pseudogenes are transposed duplications

The classification of pseudogenes was based on two criteria: i) the intron-exon structure of the pseudogene inferred from the pater gene model; ii) the relative genomic positions of pseudogene-pater pairs (Figure 1). For this analysis, a filtered dataset of 162 pseudogene-pater pairs was considered. These included pseudogenes identified by a single pater locus and paters identifying only one pseudogene. These pairs (hereafter referred to as 1:1) have minimal uncertainty in the identification of the pater locus (Supplementary Table S3). All classifications of the 1:1 dataset were subjected to manual curation (see Supplementary File 1).

**Figure 1.**
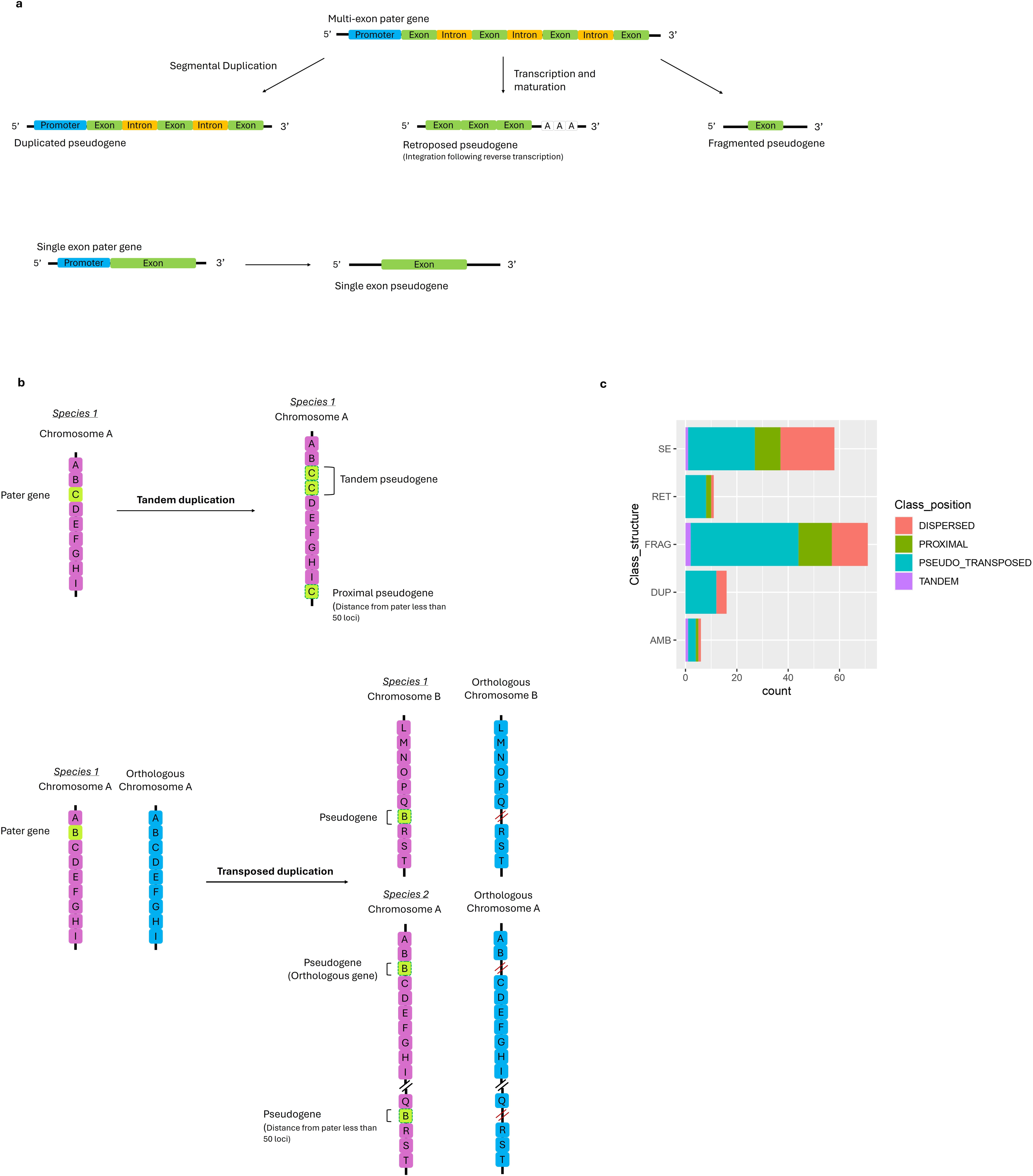
Classification of *F. graminearum* pseudogenes based on intron structure and pseudogene and pater map positions. a) Classification criterion based on intron-exon structure. b) Classification criterion based on pseudogene and pater genomic position c) Frequency distribution of 1:1 pseudogenes.

When introns were predicted in the pseudogenes at positions expected based on the pater locus model, the pseudogenes were classified as non-processed (or DUPlicated) (Figure 1a and Supplementary Table S3). The absence of introns in the pseudogenes, although expected based on the pater locus model, indicated retroposition and the pseudogene was classified as processed (or RETropseudogene) (Figure 1a). Pseudogenes showing both evidence of duplication and retroposition were classified as ambiguous (AMB) signifying uncertain classification. In cases where the alignment did not encompass intron regions, the pseudogenes were classified as fragmented (FRAG) if the pater locus had a multi-exon model, or as single exon (SE) if the pater had a model with a single coding exon (Figure 1a). The occurrence of adenine/thymidine-rich regions in close proximity of pseudogenes was examined using a sliding window approach. The presence of these regions did not provide a clear discrimination between processed and non-processed pseudogenes and thus was not adopted as an additional criterion for the classification of SE pseudogenes.

As illustrated in Figure 1c, the majority of pseudogenes (43.8%) were classified as fragmented (71 out of 162). On average, these pseudogenes were aligned to the corresponding functional loci for 14% of their coding sequence length (SE = 0.12). The second most frequent class of pseudogenes was that of single-exons, which accounted for 58 out of 162 cases (35.8%). On average, these were aligned to 30% of the functional cds length (supplemental file S1). The frequency of processed and non-processed pseudogenes was found to be similar (11 and 16, corresponding to 6.8% and 9.3%, respectively; Figure 1c and Tab. S3). On average, these were aligned to 27% and 30% of the pater locus sequence length. Only 6 pseudogenes (2.5%) had ambiguous classification (Figure 1b and supplemental file S1; Tab. S3).

Most of the pseudogene-pater pairs (91 out of 162) were classified as transposed (Figure1b-c). In all of these cases, the functional pater was classified as collinear, thereby inferring that the pseudogene was the result of a transposed duplication. Only four pseudogenes were found to be adjacent to their respective paters suggesting the occurrence of tandem duplications (Tab.S3; Figure1b-c). In contrast, 26 were separated from their paters by less than 50 loci. Thus, the underlying duplication was classified as ‘proximal’. In the remaining 40 cases, it was not possible to reconstruct a clear duplication mode, and thus the pseudogenes were classified as dispersed. No instances of collinearity were observed between the pseudogene and its corresponding pater in any of the 1:1 pseudogene-pater pairs. The inferred pseudogene structure (DUP, RET, AMB, SE.) and the pseudogene-pater duplication routes were found to be independent (P>0.05).

### Some pseudogenes are *hosted* within functional loci

Of the 234 pseudogenes that were mapped within intergenes, the median distance between the pseudogenes and the nearest genes was of 461,5 bp. A total of 144 pseudogenes were mapped within the genic model of a functional locus and were designated as ‘hosted pseudogenes’. Symmetrically, the loci with which they overlap were referred to as ‘host loci’ (see Supplementary Table S1 and Figure2). Twenty-three hosted pseudogenes were mapped within introns and 53 and 68 were mapped within the 5’ and 3’ untranslated regions (UTRs), respectively. These pseudogenic sequences may be transcribed as part of the host primary transcript. Indeed, when the pseudogene is oriented in a direction opposite to that of the host locus (*i.e*., when the host pseudogene is oriented in a forward direction while the host locus is oriented in a reverse direction, or *vice versa*), there is the possibility that host pater primary transcripts will anneal in correspondence to the pseudogene region. A total of 41 hosted pseudogenes were found to be oriented in a direction opposite to that of the pater gene.

Seven pseudogenes were mapped between a telomere and a functional locus, six of which matched the FG1G15090 pater locus. A total of 51 pseudogenes overlapped non-coding genic features, either a UTR or an intron (Tab. S2.).

### Pseudogenization plays a role in the evolution of genomic compartments

One notable feature of the *F. graminearum* genome is the presence of genomic compartments, which are distinguished by compositional and structural features, including the frequency of duplicated sequences and the number of functional loci (Zhao et al., 2014). In each genomic region, the density of pseudogene was calculated as the ratio between the number of pseudogenes and the total number of features (pseudogenes + functional loci). Genomic compartments showed significantly different pseudogene density (P<0.01). Pseudogenes were more frequent in the fast-evolving compartment than in other compartments. This observation was corroborated for compartment definitions based on collinearity analysis of the YL genome and *F. fujikoroi* (2% in the slow versus 3.5% in the fast compartment; Wilcoxon test P < 0.01) or *F. oxysporum (*1.5% in the slow versus 2.99% in the fast compartment; *Wilcoxon test P < 0.01)* (see Figure S2).

Thirty-two genomic regions with compositional indexes suggestive of RIP were identified based on the RIPPER software (van Wyk et al., 2019). Of these, 12 regions overlapped with pseudogenes. However, all regions exhibiting RIP also overlapped with repetitive regions of *F. graminearum* (supplemental file 2).

In several species, the number of splicing variants of a gene was found to be significantly correlated with the number of pseudogenes it generates (Zhang et al., 2014). In *F. graminearum*, however, this association was not statistically significant (supplemental Table S1).

Furthermore, the expression levels and patterns of the pater genes were analysed by querying the YL expression Atlas (Lu et al., 2022). Six pater loci showed no detectable expression. On average, pater loci showed a lower expression level (24,.28 *versus* 51.92) and a lower expression breadth (*i.e*., the number of tissues in which a gene is expressed; 7.22 *versus* 7,62) than non-pater loci (Wilcoxon test P < 0.01; Tab. S1). Finally, an examination of the gene ontology of the pater loci revealed several functions linked to pathogenesis (Figure 3), including 3,4 hydrocoumarine hydrolase activity (Deryabin et al., 2021), oxophytodienoate reductase (Degtyaryov et al., 2023).

**Figure 2.**
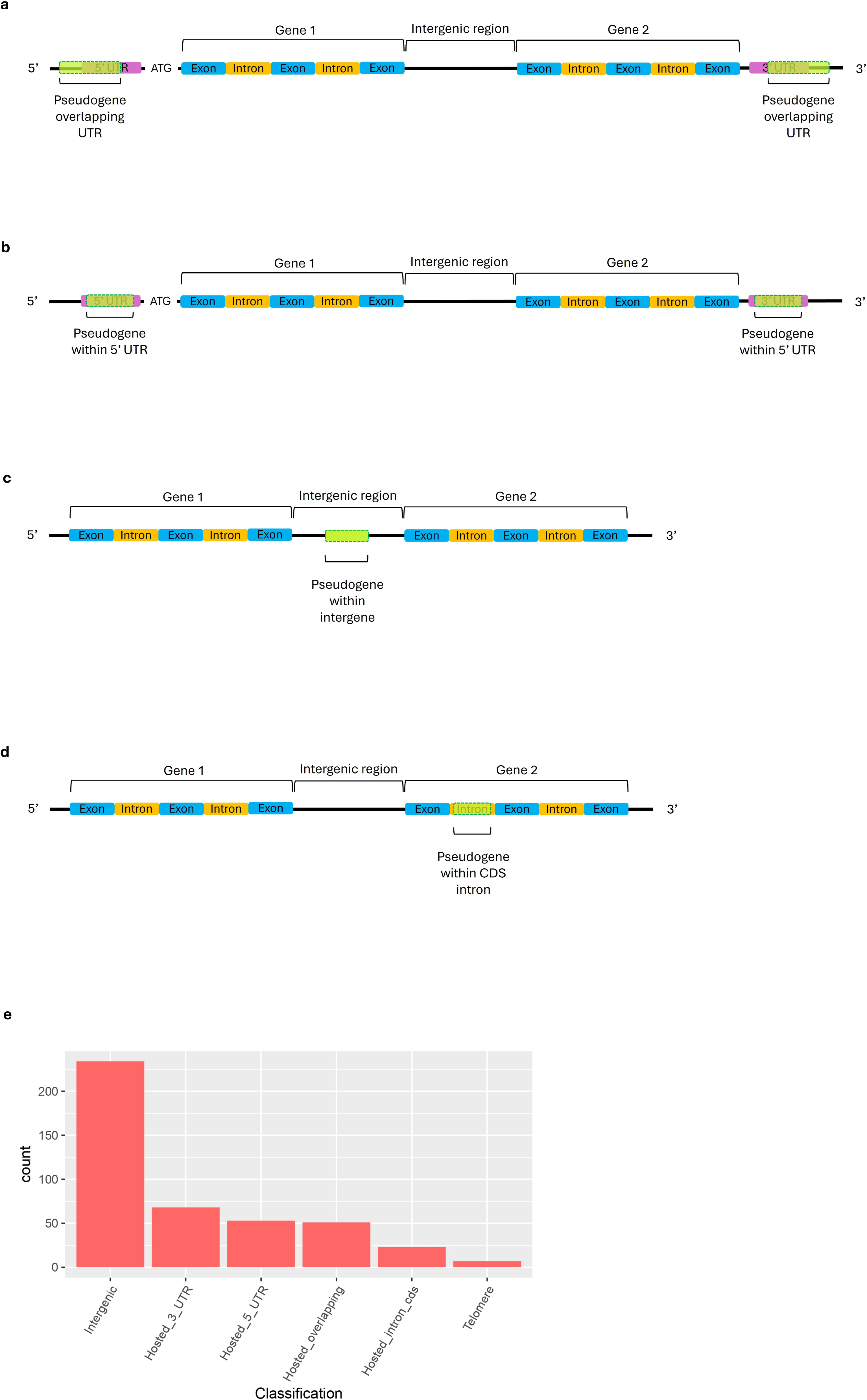
Classification and distribution of 1:1 pseudogenes. a) Hosted pseudogenes overlapping the 5’ and 3’ UTRs, b) Hosted pseudogenes mapped within 5’ and 3’ UTRs, c) Pseudogenes mapped within intergenic regions, d) Hosted pseudogenes mapped within introns, e) Distribution of *F. graminearum* pseudogenes according to their genomic positions.

**Figure 3.**
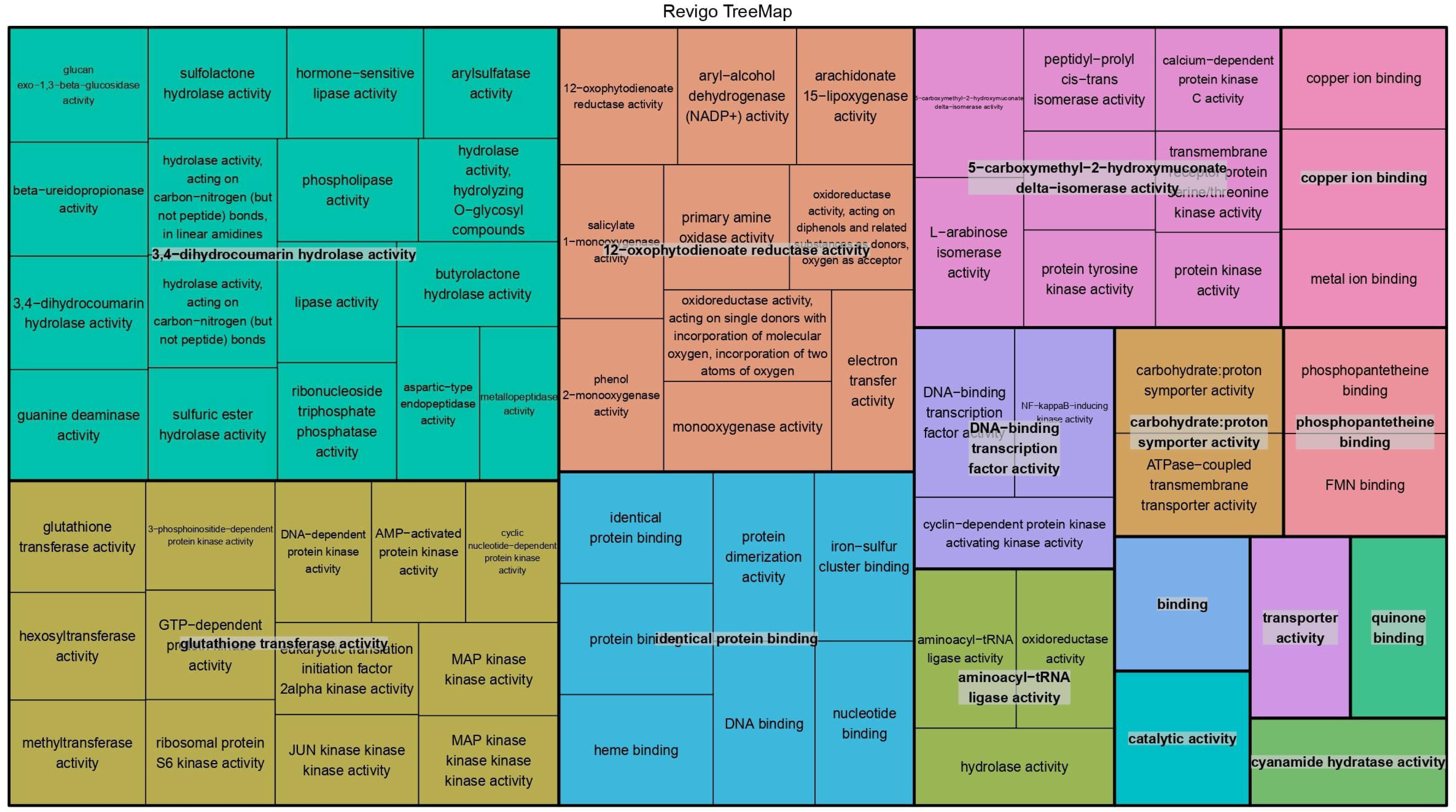
TreeMap of GO terms of paters.

### Unitary pseudogenes

A total of 544 unitary pseudogenes were identified by protein-coding loci from five *Fusarium* species (*F. culmorum*, *F. fujikoroi*, *F. oxysporum*, *F. verticillioides*, *F. solani*) (Tab. S4). As all the query pater loci were selected due to the absence of a functional orthologous locus in the YL genome, the identified YL matching sequences could be considered as putative cases of loss of functions (LOF) (Zhang et al., 2010).

A total of 203 unitary pseudogenes were collinear with their respective pater loci (Tab. S5).

These pseudogenes are extant, inactive relics of functional genes. We hypothesized that they originally contained the same repertoire of functional domains as their orthologous paters. The paters of 18 LOF pseudogenes showed homology to at least one PFAM domain. We found several domains previously identified in proteins involved in pathogenesis in *Fusarium* spp. For example, the pseudogene identified by FOXG_12385 is a *Fusarium oxysporum* locus encoding a protein with two domains, PF05368.13 and PF13460.16 domains, that were both found in the NmRA transcription factors. The NmrA transcription factors participate in the nitrogen-responsive pathway which operates via TOR to control virulence in several pathogenic fungi (Pfannmüller et al., 2017; Teichert et al., 2006). It is noteworthy that FOXG_12385 displays orthologs In *F. fujikoroy*, *F. solani,* and *F. culmorum* but not in *F. verticilliodes* and *F. graminearum.* Another noteworthy example is that of the pseudogene which matches FOXG_07976 a protein with the GLEYA domain which is typical of fungal adhesins (Gámez-Arjona et al., 2022). Again, the pseudogene identified by FOXG_07068T0 encodes a protein with the PF12013.8 domain which is known to be present in orsellinic acid/F9775 biosynthesis cluster proteins. Orsellinic acid is a simple aromatic poliketyde formed through the condensation of acetyl-CoA and malonyl-CoA and is the precursor of a wide range of derivatives (Jørgensen et al., 2014; Tao and Abe, 2021) Noteworthy, *F. graminearum* contains 15 polyketide synthase (PKS) genes, one of which, PKS14, is involved in orsellinic acid and orcinol synthesis whin has been shown to be expressed during plant infections or when cultivated on rice or corn meal based media (Jørgensen et al., 2014).

## Discussion

The release of the *Fusarium* genome version YL has provided the scientific community with a highly accurate annotation of genomic features (Lu et al., 2022). Such a dataset is optimal for the identification of pseudogenes through approaches based on sequence homology to functional genes. Through the employment of Exonerate as an alignment tool, we have identified 1862 *F. graminearum* non-coding sequences that show homology to paralogous functional proteins. These regions were further characterised in relation to the structural features of matching functional genes, thus defining the *F. graminearum* pseudogene complement.

### Genomic distribution of pseudogenes in *F. graminearum*

The pseudogenes that are identified by *F. graminearum* proteins are distributed across all four chromosomes with a notable cluster present in the subtelomeric region of chromosome 4. This cluster corresponds to seven functional loci, all of which are situated near one another and represent an array of tandem duplications. Given the similarity in structure observed among most pseudogenes within this cluster, we postulate that the duplications in question represent inactivated copies.

Apart from this region no significant differences in pseudogenes density were observed between chromosomes. However, it was noted that the pseudogenes tend to accumulate preferentially in the fast-evolving genomic compartment of each chromosome. Furthermore, our data on pseudogene characterization indicates that up to 91 of the gene copies were inactivated following transposition. This finding may be indicative of either a higher rate of sequence turnover in these regions and/or a tendency for sequence duplication by transposition at specific sites. Indeed, based on the evidence that *F graminearum* relocated genes map preferentially to these regions, Zhao et al. (2014) have proposed that these sites are source of gene innovation and recombination (Croll and McDonald, 2012).

### Is there a preferred mechanism for sequence duplication in F. graminearum?

No significant associations were identified between the classifications of pseudogenes based on map positions and those based on their inferred structure. The data collectively indicate that there is no preferential mechanism for sequence duplication in *F. graminearum*.

Studies in other systems have shown that the formation of pseudogenes occurs through a variety of mechanisms. For example, in mammals, pseudogenes are preferentially generated by retroposition (Sisu et al., 2014; Zhang and Gerstein, 2004; Zheng et al., 2007), whereas in the fly they are mostly generated by segmental duplication (Sisu et al. 2014). The situation in plants is more complex, with several lines of evidence indicating that the majority of pseudogenes are classified as fragmented or single exon and are likely to arise by a mechanism of duplication by transposition involving double-strand break repair (Mascagni et al., 2021; Prade et al., 2018; Xie et al., 2019; Zou et al., 2009) Following the findings in plants and considering the abundance of fragmented pseudogenes in *F. graminearum,* it can therefore be postulated that the enrichment of pseudogenes in the fast-evolving compartment reflects a higher frequency of double-strand breaks or the type of repair system active at these sites. An alternative hypothesis is that there is a higher frequency of duplications in the fast-evolving regions, which are then converted into pseudogenes by the Repeat Induced Point (RIP) mutation mechanism, in accordance with the observation that RIP is active in *F. graminearum* No clear associations between pseudogenes and *in silic*o-identified RIPPED regions were found. Further studies are therefore required to ascertain whether the sequence composition of pseudogenes is due to mutational bias active in the regions where they reside, or alternatively, whether they are the result of RIP events that have been overlooked by our *in silic*o approaches. It has been hypothesised that a direct consequence of RIP in *F. graminearu*m is the low frequency of duplications in contrast to the high frequency of singleton genes (Cuomo et al., 2007; King et al., 2015). The relationship between pseudogenes and gene dosage lends support to two alternative hypotheses. One might posit that pseudogenes should be preferentially associated with pater genes that are members of gene families. Indeed, the presence of multiple copies of genes is associated with a relaxed purifying selection, which in turn increases the probability of inactivation (Chang et al., 2020; Panchy et al., 2016). Furthermore, the existence of families of homologous genes is in accordance with the existence of RIP escape/avoidance features that may be conserved also in the pseudogenes (Panchy et al., 2016). The second hypothesis is that pseudogenes are records of gene decay events that should regulate copy number. Consequently, one might expect that most of the paters are actually singletons because their copies have become pseudogenes (Cuomo et al., 2007). However, no significant association was found between the propensity to give rise to pseudogenes and the duplication depth of the paters .

### Unitary genes, loss of function and evolution of pathogenicity and virulence

It is established that the genomes of *Fusarium species* contain a considerable number of species-specific genes, which are thought to have been acquired by horizontal transfer processes (Cuomo et al., 2007). The presence of horizontally inherited genes may represent a crucial step in the evolution of pathogenicity and virulence. Pseudogenes may prove to be a source of valuable information in this context, as they represent a record of the opposite phenomenon: the loss of functions. A number of compelling examples illustrate the contribution of loss function through pseudogenization to the virulence of pathogenic microorganisms. This phenomenon can be exemplified by the observations of Yang et al. (2024) (Yang et al., 2024), who noted that the loss of a gene can induce a fitness advantage for the strains, thereby enabling them to adapt to the surrounding environment. To give an example, the pseudogenisation of several functional genes in Salmonella has resulted in a variation in tolerance to ROS (such as HlJOlJ), an increased pathogenicity due to an enhanced ability to damage epithelial cells, a higher capability to produce systemic infections, and an impact on flagella formation. The latter not only endows bacteria with the capacity for mobility but also plays a pivotal role in the processes of adhesion, invasion and biofilm formation, which are integral to the development of bacterial pathogenicity. A recent comparative genomic study in fungi has revealed that the evolutionary route from a unicellular opisthokont ancestor to derived multicellular fungi involved a combination of gradual gene loss and several large-scale duplication events. These findings challenge the traditional view that abrupt changes are the primary drivers of fungal evolution. Instead, they suggest that an interplay of gene gain and loss plays a significant role in shaping fungal evolution (Merényi et al., 2023) .

In our study, we investigated the phenomenon of loss if function by analysing the repertoire of unitary pseudogenes in several Fusarium species. These are genic-like sequences that do not correspond to functional paralogous; instead, they correspond to functional orthologous sequences. The protein sequences from five *Fusarium* species were used to query the YL genome, which had been masked for repetitive, coding sequences and pseudogenic regions matching paralogous loci. A significant proportion of orthologous pater sequences were collinear with the pseudogenes, suggesting that they represent records of inactivated *F. graminearum* singletons. The distribution of pseudogenes between genomic compartments showed an enrichment for the fast-evolving sites, confirming the heightened evolutionary dynamism of these regions. Most unitary paters of *F. culmorum* did not present orthologs in the other investigated species. These instances merit scrutiny. To minimise the impact of artefacts resulting from the *ab-initio* annotation of the orthologous genome we focused our analysis on the paters that exhibited evidence of functional domains. The application of this filter resulted in a significant reduction in the number of identified pater pseudogenes. The utilisation of long-read sequencing for the analysis of the transcriptome and the refinement of gene predictions will facilitate more comprehensive characterization of unitary paters, thereby enabling a more detailed investigation into the extent to which gene loss contributes to the evolution of *F. graminearum* and other Fusarium species or to the dynamics of pathogenicity and virulence.

The list of pseudogenes compiled in this study is a valuable starting point for future research aimed at elucidating the genetic bases of environmental adaptation and bridging the gap between genetic and phenotypic architecture in Fusarium species. The pseudogenes identified in our study can be considered as interesting potential candidates, especially for their involvement in life history traits and other relevant virulence-related traits. This is suggested, for example, by the observation that the paters of 18 loss-of-function pseudogenes showed homology to domains previously identified in proteins involved in pathogenesis, such as PF05368.13 and PF13460.16, both found in the NmRA transcription factors, GLEYA, typical of fungal adhesins, and PF12013.8, known to be present in the orsellinic acid/F9775 biosynthesis cluster proteins. Furthermore, is noteworthy that approximately one-third (144/436) of the pseudogenes were found to overlap with untranslated or intron sequences of functional loci, indicating the potential to be transcribed. Very intriguingly, Qi et al. (2024) demonstrated empirically, working with 71 investigated *Fusarium graminearum* pseudogenes, that a mechanism known as ‘restorative RNA editing’ can confer adaptive advantages in fungi to resolve survival-reproduction trade-offs caused by antagonistic pleiotropy and confer a selective advantage in fungi (Qi et al., 2024).

Indeed, in fungi, restorative editing corrects premature stop codons in pseudogenes to their ancestral state specifically during sexual reproduction. Qi et al. (2024) have shown that 16 pseudogenes undergoing restorative editing are critical for the development of germline tissues in fruiting bodies, and have also observed that i) restorative RNA editing facilitates the emergence of premature stop codons, ii) that the stop codons corrected by restorative editing are favoured over the ancestral codons, and iii) that the ancestral versions of the pseudogenes have antagonistic effects on reproduction and survival. Thus, the present study paves the way for studies aimed at identifying other pseudogenes that may contribute to the fitness of *F. graminearum*.

## Experimental procedures

### Sequence dataset

All genomic sequences and annotations used in this study were obtained from public databases. For *F. graminearum,* we used the genomic sequences of the PH1 strain and the transcript-annotation 5’collapsed downloaded from http://fgbase.wheatscab.com/. For *F. culmorum*, we used the assembly of chromosomes 1-4 deposited as assembly GCA_900074845.1 (assembly name FcUK99v1.2). The other contigs were not considered as they are assembled without physical map evidence. For *F. fujikoroi,* the genome-build accession GCA_000315255.1 from the Joint Genome Institute EF 1 (genome version EF1) was used. The genome sequence, assembly, and annotation of protein-encoding genes of the *F. oxysporum* genome were generated by the Broad Institute as part of their work on the Fusarium Comparative Sequencing Project (genome-build GCA_000222805.1, genome version PO2). The assembly and annotation of *F. solani* used in this study were obtained for JGI (V2.0 INSDC assembly GCA_000151355.1). The assembly and annotation of *F. verticillioides* were provided by the Broad Institute (genome build GCA_000149555.1). Exons, introns, and intergenic sequences were extracted from the genome sequence using the gff2sequence software (Camiolo and Porceddu, 2013).

### Identification of pseudogenes

Pseudogenes were identified by homology search using Exonerate software (Slater and Birney, 2005) considering YL proteins as query and masked YL genome as subject. All YL proteins were used to identify pseudogenes with functional paralogous. The subject genome was masked in correspondence of repeats and coding sequences using Repeatmasker (Smit et al., 2013). Pairs of hits that overlapped by more than 20% of the length of the shortest hit were included in a cluster featuring a pseudogene region. For each of these clusters the pseudogene-query pair with the highest Exonerate score and alignment identity was selected. The selected functional query was considered to be the pater locus, and its structure was assumed to be the best existing approximation of the pseudogene structure at the time of its origin (Zhang et al., 2006).

Orthologous groups in *F. graminearum*, *F. culmorum*, *F. fujikoroi*, *F. oxysporum*, *F. solani* and *F. verticilliodes* were identified using OrthoFinder with default settings (Emms and Kelly, 2019). Unitary pseudogenes in the YL genome were identified with exonerate (Slater and Birney, 2005), using as queries the proteins encoded by loci with no ortholog in *F*. *graminearum*. The subject genome sequence was masked in regions corresponding to repeat elements, annotated functional loci and previously identified pseudogenes. Hits identified by Exonerate were processed as described above (see Figure S1). To identify A or T (A|T) rich regions in the YL genome, we applied a sliding window approach to the YL genomic sequence. An A|T rich region of size n contained at least k A or T nucleotides (with k<n). A|T rich region was considered a putative site of polyA integration during retroposition if its distance from a pseudogene was less than 600 bp.

### Classification of pseudogenes based on intron-exon structure

The structure of a pseudogene was defined considering the exonerate output and the Genewise alignment of the pater’s protein to the pseudogene sequence (Slater and Birney, 2005) Genewise was run with default settings to align the pater protein to the genomic sequence of the pseudogene and to infer the intron-exon structure. The pseudo-exon-intron structure was then compared with the YL pater gene model to check for the presence of matching intron-exon junctions.

Non-processed pseudogenes originate from chromosomal or segmental duplications and are therefore expected to retain the intron-exon structure of the pater loci. The pseudogene exon-intron junctions that matched the position of the corresponding junction in the pater locus were classified as DUP, which stands for Duplicated (Mascagni et al., 2021) (Figure 1). Exon-intron positions showing stretches of gaps in the pater protein, although no intron was predicted by Genewise, were also classified as ‘DUP. The opposite situation, *i.e*., gaps in the pseudogene where an intron is expected based on the gene structure of the pater locus, were coded as ‘AMB (Ambiguous), to indicate uncertainty. Cases where exonerate indicated the presence of two pseudoexons but Genewise was unable to identify an exon-intron junction in the pseudogene, were classified as ‘dup’ if the predicted pseudoexons were more than 50 bp apart. Alignment positions that covered an intron-exon junction in the pater structure and had no inferred introns by Genewise were classified as ‘ret’ (=retroposed), if the pseudogene sequences showed no homology to the pater intron sequence in Blastn searches.

Pseudogene models with all exon-intron junctions classified as ‘dup’ and none as ‘ret’ were considered as non-processed or duplicated. Those with ‘RET’ positions and non as ‘DUP or ‘AMB were classified as processed or retroposed. Pseudogenes identified by single exon paters were classified as ‘single exon’ pseudogenes, whereas those identified by multi-exon paters but whose alignment did not cover the intron-exon junctions of the pater locus were classified as ‘fragmented’ (Figure S1). Finally, pseudogenes with ‘AMB or containing both ‘DUP and ‘RET exon-intron junctions were classified as ambiguous.

The polyA tails of pseudogenes, a hallmark of retroposition, were identified as genomic regions with more than a minimum number of consecutive adenines (16) tolerating at most two mismatches. Finally, the software tfasty36 was used to identify disablements (frameshifts or stop codons) in the sequence alignments of the pater protein to the pseudo-exon sequence.

### Classification of pseudogenes according to duplication mode

Sequence duplicates were classified using McScanX (Wang et al., 2013). Briefly, each query protein of a species was used as a subject in pairwise Blastp searches against the entire proteome. The top five non-self-matching hits with an e-value below 1x e^-10^ were filtered for further analysis. The hits were classified according to their position in the genome. If the loci of a hit were collinear, the duplication was classified as segmental (and coded as ‘wgd’). Hits identified by paters mapping to the same chromosome were classified as tandem duplications if the pseudogene and the pater were adjacent, and otherwise as proximal if they were less than 50 non-homologous loci apart. The remaining pseudogene-pater pairs were identified as transposed if one locus of the pair had a collinear ortholog in another species, and otherwise as dispersed.

### Gene ontology and protein domain family enrichment analysis

Pseudogenes were predicted to have the same domain as their paters and were therefore referred to the same GO terms as their paters. GO terms were summarised with REVIGO (Supek et al., 2011). GO enrichment tests were performed with TopGO using the ‘elim’ algorithm.

## Acknowledgements

The authors are indebted with the FAR for financial support. The authors have no conflict of interest.

