## Supplementary figures and images for "Identification and structural characterization of pseudogenes in *Fusarium graminearum*"

### Supplemental Figure 1

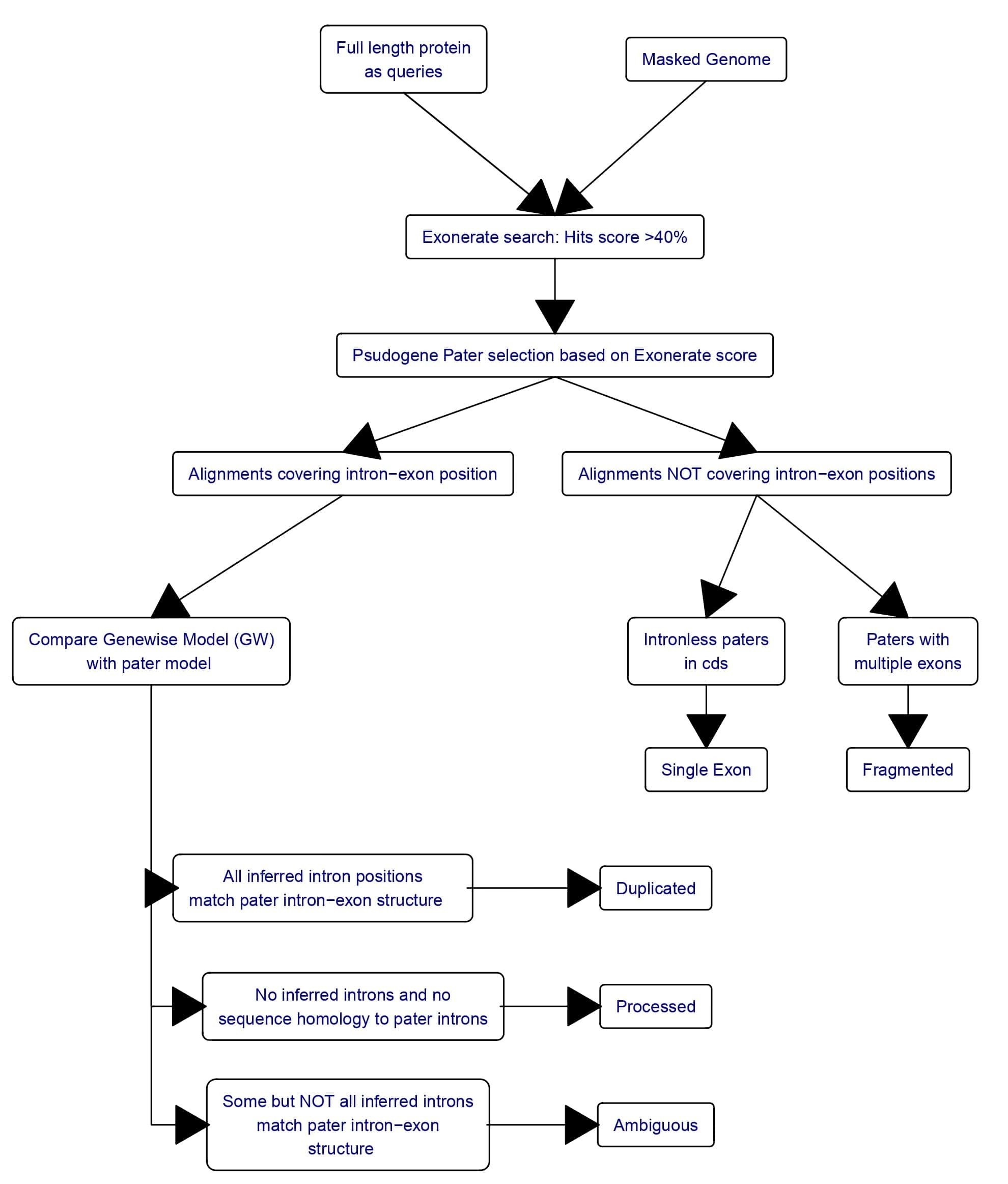
